# Polymerization Kinetics Stability, Volumetric Changes, Apatite Precipitation, Strontium Release and Fatigue of Novel Bone Composites for Vertebroplasty

**DOI:** 10.1101/468652

**Authors:** Piyaphong Panpisut, Kirsty Main, Muhammad Adnan Khan, Mayda Arshad, Wendy Xia, Haralampos Petridis, Anne Margaret Young

## Abstract

**Purpose:** The aim was to determine effects of diluent monomer and monocalcium phosphate monohydrate (MCPM) on polymerization kinetics and volumetric stability, apatite precipitation, strontium release and fatigue of novel dual-paste composites for vertebroplasty.

**Materials and methods:** Polypropylene (PPGDMA) or triethylene (TEGDMA) glycol dimethacrylates (25 wt%) diluents were combined with urethane dimethacrylate (70 wt%) and hydroxyethyl methacrylate (5 wt%). 70 wt% filler containing glass particles, glass fibers (20 wt%) and polylysine (5 wt%) was added. Benzoyl peroxide and MCPM (10 or 20 wt%) or N-tolyglycine glycidyl methacrylate and tristrontium phosphate (15 wt%) were included to give initiator or activator pastes. Commercial PMMA (Simplex) and bone composite (Cortoss) were used for comparison.

ATR-FTIR was used to determine thermal activated polymerization kinetics of initiator pastes at 50-80 °C. Paste stability, following storage at 4-37 °C, was assessed visually or through mixed paste polymerization kinetics at 25 °C. Polymerization shrinkage and heat generation were calculated from final monomer conversions. Subsequent expansion and surface apatite precipitation in simulated body fluid (SBF) were assessed gravimetrically and via SEM. Strontium release into water was assessed using ICP-MS. Biaxial flexural strength (BFS) and fatigue properties were determined at 37 °C after 4 weeks in SBF.

**Results:** Polymerization profiles all exhibited an inhibition time before polymerization as predicted by free radical polymerization mechanisms. Initiator paste inhibition times and maximum reaction rates were described well by Arrhenius plots. Plot extrapolation, however, underestimated lower temperature paste stability. Replacement of TEGDMA by PPGDMA, enhanced paste stability, final monomer conversion, water-sorption induced expansion and strontium release but reduced polymerisation shrinkage and heat generation. Increasing MCPM level enhanced volume expansion, surface apatite precipitation and strontium release. Although the experimental composite flexural strengths were lower compared to those of commercially available Simplex, the extrapolated low load fatigue lives of all materials were comparable.

**Conclusions:** Increased inhibition times at high temperature give longer predicted shelf-life whilst stability of mixed paste inhibition times is important for consistent clinical application. Increased volumetric stability, strontium release and apatite formation should encourage bone integration. Replacing TEGDMA by PPGDMA and increasing MCPM could therefore increase suitability of the above novel bone
composites for vertebroplasty. Long fatigue lives of the composites may also ensure long-term durability of the materials.

## 1. Introduction

Osteoporotic fracture of the spine (osteoporotic vertebral fracture; OVF) causes severe pain, height loss, limited mobility, kyphosis, and reduced pulmonary function [1]. Non-surgical treatments such as analgesics and rehabilitation are commonly used but often fail to relieve severe pain in some patients [2, 3]. Hence, surgical managements that relieve severe pain rapidly such as vertebroplasty (VP) and balloon kyphoplasty (KP) are indicated. These procedures involve injection of a bone cement to stabilize fractures. Common complications of these treatments are cement leakage (up to 77 %) leading to neurological deficits [4], adjacent vertebral fractures (12 - 15 %) [5], and post-operative infection which can be a rare but serious complication [6].

Polymethyl methacrylate (PMMA) cement is the most commonly used bone cement for VP and KP. Limitations of this cement include poor controlled setting and viscosity that may increase the risk of cement leakage [7]. Further concerns are high polymerization shrinkage, heat generation, and risk of toxic unreacted monomers release [8]. These shortcomings may cause gap formation and local inflammation leading to fibrous encapsulation [9] reducing the integrity of the bone-cement interface. Furthermore, conventional PMMA cements also lack the ability to promote bone formation.

Two-paste injectable bone composites have been developed to address some limitations of the PMMA cements but various shortcomings remain. For example, the primary base monomer used has been bisphenol A-glycidyl methacrylate (Bis-GMA).

This monomer is known to limit final monomer conversion of composites due to its limited mobility [10]. Additionally, the commercial composites contain TEGDMA as a diluent monomer, which is known to increase shrinkage and heat generation of dental composites due to its high density of methacrylate groups [10, 11]. Furthermore, the composites contain the tertiary amine DMPT (N,N-dimethyl-p-toluidine), which is highly toxic to human cells [10, 12]. Recently developed light-activated urethane dimethacrylate (UDMA)-based dental composites exhibited higher monomer conversion than Bis-GMA based commercial composites [10]. The same study also demonstrated that replacing TEGDMA by polypropylene glycol dimethacrylate (PPGDMA) increased monomer conversion and cytocompatibility of dental composites while polymerization shrinkage was reduced.

Following mixing, chemically-activated bone composites should ideally cure rapidly after a well-defined inhibition time that provides sufficient working time for injection. A potential problem, however, is unmixed paste instability due to thermal initiated polymerization arising upon storage [13]. Manufacturers usually recommend chemically-activated paste storage below 4 °C [14]. This ideal storage condition, however, may be difficult to achieve in some circumstances. For example, medical products shipping to tropical regions can expose materials to fluctuating temperatures between - 4 to 42 °C and 10 to 40 °C during air and marine transport respectively [15]. Stability may be estimated from inhibition times at elevated temperatures for unmixed pastes. Temperature dependence of these times is expected to be governed by Arrhenius type equations, be directly proportional to concentration of initiator (benzoyl peroxide, BP) and inversely proportional to inhibitor levels added to stabilise different monomers [16].

The addition of monocalcium phosphate (MCPM) and tri calcium/strontium phosphates (TCP / TSrP) into dental composites has been shown to promote hygroscopic expansion which could potentially balance polymerisation shrinkage and relieve residual shrinkage stress [17, 18]. The addition of these reactive phosphates also enabled surface apatite formation which is known to correlate with *in vivo* bone bonding [19, 20]. Apatite formation can also be enhanced by the addition of polylysine (PLS) [17]. Furthermore, strontium can promote osteoblast proliferation and maturation whilst inhibiting osteoclast activities [21-23]. It will also enhance radiopacity [24], which may facilitate the surgical procedure and enable follow up with radiographs.

Injected bone composites should be able to withstand the fluctuating and repetitive loads during physical activities [25]. This may then help to prevent mechanical failure due to crack propagation induced by repetitive subcritical loads (fatigue failure). A recent study [26] indicated that high strength values of composites displayed under static loading were not directly related to fatigue performance. Fatigue of various materials was previously assessed by generating stress versus number of cycles curve (S-N curve) during simulated fatigue [27, 28]. At a given applied stress, the steep gradient of *S-N* plot was associated with a significant reduction in failure cycles [29]. Therefore, a low gradient rather than high gradient was preferable in terms of fatigue performance [30].

The aim of this study was to compare TEGDMA/UDMA versus PPGDMA/UDMA-based bone composites with added Ca/Sr phosphates (MCPM and TSrP) and polylysine (PLS). Initiator paste stability, kinetics of polymerization, final monomer conversions, polymerization shrinkage and heat generation, water sorption induced mass and volume changes, surface apatite formation, strontium release, and biaxial flexural strength / fatigue were assessed. The effect of MCPM levels (5 wt% versus 10 wt%) and type of diluent monomers (TEGDMA versus PPGDMA) were examined.

## 2. Materials and Methods

### 2.1 Material paste preparation

Experimental bone composites were prepared using a powder to liquid mass ratio of 70: 30. The liquid phase (Table 1) contained urethane dimethacrylate (UDMA) (MW 479 g/mol, DMG, Hamburg, Germany), polypropylene glycol dimethacrylate (PPGDMA) (MW 600 g/mol, Polyscience, PA, USA) or triethylene glycol dimethacrylate (TEGDMA) (MW 286 g/mol, DMG, Hamburg, Germany), and hydroxyethyl methacrylate (HEMA, MW 130 g/mol) (DMG, Hamburg, Germany). To this was added either benzoyl peroxide (BP) (MW 242 g/mol Polyscience, PA, USA) for the initiator liquid or N-tolyglycine glycidyl methacrylate (NTGGMA) (MW 329 g/mol, Esschem, Seaham, UK) for the activator liquid.

**Table 1.** Components of liquid phases before mixing with the powder phase. Upon mixing the composite, BP and NTGGMA concentrations will become 1.5 and 1 wt% respectively.

| Liquid phase / components | UDMA | PPGDMA /TEGDM | HEMA | BP | NTGGMA |
| --- | --- | --- | --- | --- | --- |
|  | wt% of monomers | wt% of monomers | wt% of monomers | wt% of liquid | wt% of liquid |
| Initiator liquid | 70 | 25 | 5 | 3 | 0 |
| Activator liquid | 70 | 25 | 5 | 0 | 2 |

Powder phase (Table 2) contained glass filler (particle diameter of 0.7 μm, DMG, Hamburg, Germany), glass fiber (30 μm in diameter and 150 μm in length, Mo-Sci, PA, USA), monocalcium phosphate monohydrate (MCPM) particle diameter of 53 μm, Himed, NY, USA), tristrontium phosphate (TSrP) (particle diameter of 10 μm, Sigma Aldrich, Gillingham, UK), and polylysine (PLS) (particle diameter of 20-40 μm, Handary, Brussel, Belgium).

**Table 2.**
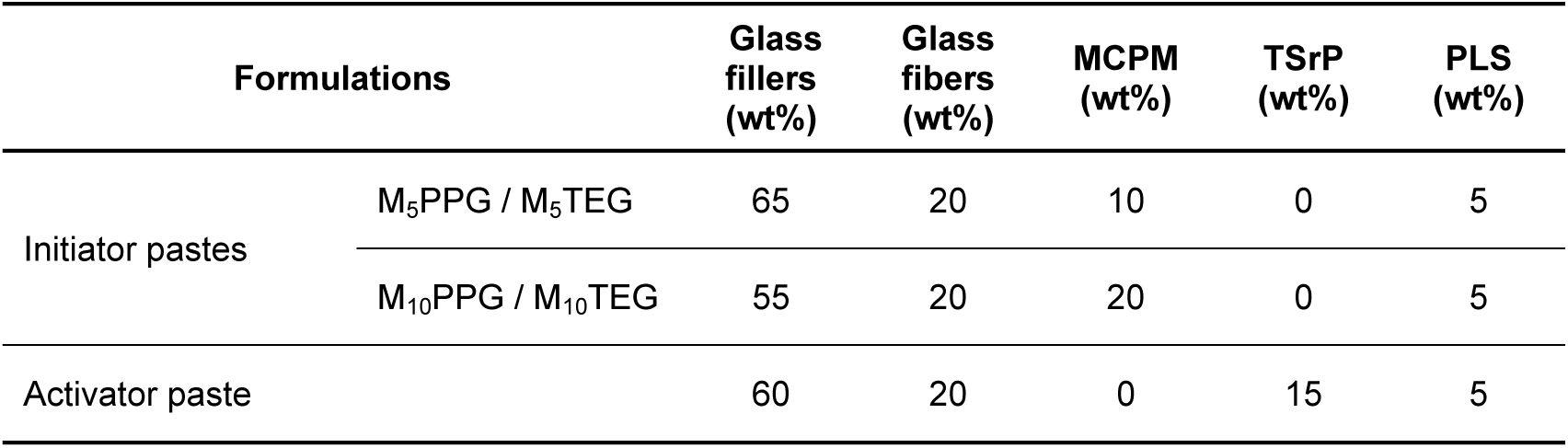
Components of powder phase. Formulations with varying level MCPM (5, 10 wt%) and types of diluent monomer (PPGDMA, TEGDMA). The powder phase of each formulation was mixed with PPGDMA (PPG) or TEGDMA (TEG) liquid phases presented in Table 1. MCPM and TSrP in filler are halved after mixing initiator and activator paste.

Composite pastes were prepared at 23 °C. Powders and monomers were weighed using a four-figure balance (OHAUS PA214, Pine Brook, USA). The powder phase was mixed with the liquid phase containing either initiator or activator using a planetary mixer (SpeedMixer, DAC 150.1 FVZ, Hauschild Engineering, Germany) at 2000 rpm for 2 min. The initiator and activator pastes were then poured into a double-barrel syringe (MIXPAC, SULZER, Switzerland) over a vibrator to reduce air entrapment. The syringe was left in an upright position for 24 hr at 23 °C to allow the release of air bubbles. For stability studies, pastes were then stored at 4, 23, and 37 °C for 1, 3, 6, 9, and 12 months to give “aged” pastes. Mixed experimental initiator and activator pastes were obtained using an automatic mixing tip attached to the double-barrel syringe and a mixing gun (MIXPAC Dispenser, SULZER, Switzerland). Commercial PMMA cement (Simplex) and bone composite (Cortoss) (Table 3), mixed as per manufacturer’s instructions within their use by date, were used as comparisons.

**Table 3.** Components of commercial products.

| Commercial materials | Compositions | Suppliers |
| --- | --- | --- |
| Simplex P®<br>(powder and liquid) | Liquid: DMPT, MMA, inhibitor<br>Powder: BP, PMMA, MMA-styrene copolymer beads (diameter of ~25 µm), barium sulphate (diameter ~ 2 µm) | Stryker,<br>Berkshire, UK |
| Cortoss®<br>(double-barrel syringe) | Liquid phase: Bis-GMA, Bis-EMA, TEGDMA, BP, DMPT, inhibitor<br>Powder phase: glass ceramic particles (combeite, diameter of ~ 5-30 µm) |  |

### 2.2 FTIR studies of composite pastes

#### 2.2.1 Monomer conversion profiles

Monomer conversion profiles of pastes (n=3) were obtained using FTIR (Perkin-Elmer Series 2000, Beaconsfield, UK) with a temperature controlled ATR attachment (3000 Series RS232, Specac Ltd., UK). Initiator pastes or mixed experimental bone composites and commercial materials were placed in a metal circlip (1 mm depth and 10 mm diameter) on the ATR diamond and covered with an acetate sheet. FTIR spectra between 700 - 4000 cm^−1^ of the bottom surfaces of the specimens were recorded every 4 s at a resolution of 4 cm^−1^. For unmixed pastes, spectra were recorded for up to 10 hours at 50, 60, 70 or 80 ± 1 °C. With mixed pastes, spectra were obtained for 40 min at 25 ± 1 °C.

Monomer conversion, D_c_, and rate of polymerization, R_p_, versus time were obtained from FTIR spectra using equations 1 and 2. [17].

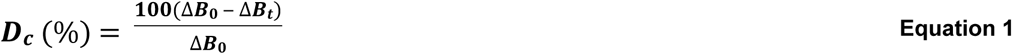

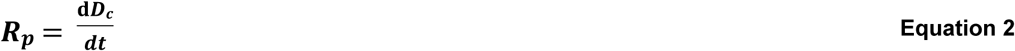

Where Δ*B*_0_ and Δ*B_t_* were the absorbance of the C-O peak (1320 cm^−1^) above background level at 1335 cm^−1^ initially and after time t and dD_c_/dt was the gradient of conversion versus time. Furthermore, final monomer conversion, D_c,max_, was calculated by linear extrapolation of conversion versus inverse time to zero.

An example of a monomer conversion and reaction rate profile is shown in Fig 1A and B. These demonstrate a delay time (inhibition time) before rapid rise in monomer conversion (snap set). Maximum rate is observed between 10-40% conversion with the reaction slowing significantly thereafter.

**Fig 1.**
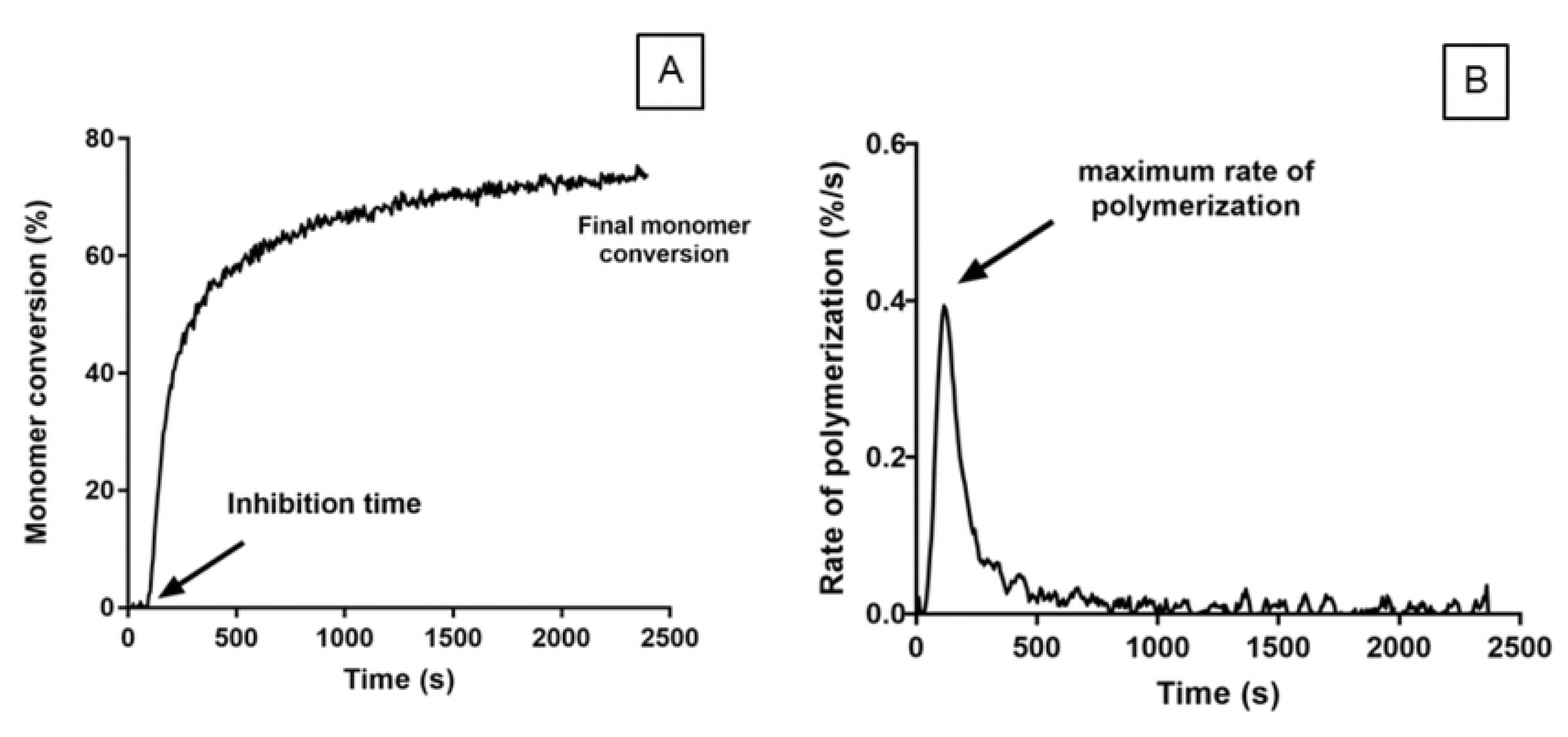
Example profiles of A) polymerization and B) rate of polymerization of mixed M_10_PPG. All mixed and unmixed pastes exhibited similar features for both profiles.

The standard mechanism of free radical polymerisation of dimethacrylate monomers includes, initiation, inhibition, propagation, crosslinking and termination steps. Using this mechanism, with the stationary state assumption that the concentration of free radicals is constant, gives the inhibition time as [16]:

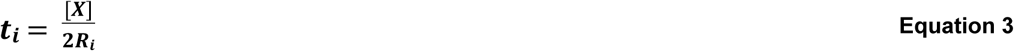

[X] is the initial concentration of inhibitor and R_i_ the rate of initiation. Furthermore, rate of polymerization (R_p_) can be described using the following equation[31].

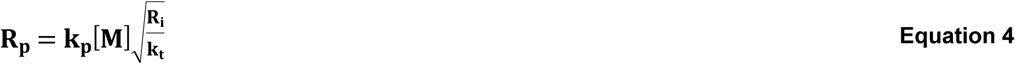

[M] is the monomer concentration and k_p_ and k_t_ rate constants for propagation and termination steps. Combining equations 3 and 4, 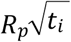 is therefore expected to be independent of rate of initiation and given by the following equation.

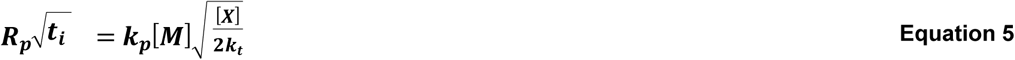

In the following, inhibition times were calculated by linear extrapolation of data between 10% and 40% monomer conversion back to 0% conversion. The gradient in this range was used to obtain the maximum rate of polymerization R_p,max_ and 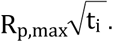

#### 2.2.2 Thermally activated polymerization of unmixed initiator paste, activation energies and predicted shelf life

Pilot studies revealed that initiator pastes were more susceptible to heat than activator pastes and that modifying MCPM level had relatively minimal effect. Hence, initiator pastes of M_10_PPG and M_10_TEG were chosen to assess polymerisation kinetics and thermally activated polymerization of the experimental bone composites. FTIR spectra of freshly mixed M_10_PPG and M_10_TEG initiator pastes (n=1) were used to obtain their inhibition times, rates of polymerization and final conversions at temperatures of 50, 60, 70, and 80 °C.

Inverse inhibition times and reaction rates are proportional to rate constants, k, whose temperature dependence, are generally described by Arrhenius type equations [32].

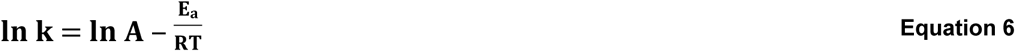

T is temperature in Kelvin and R the gas constant. A is a pre-exponential factor that is related to the frequency of molecular collisions between reacting species and E_a_, the activation energy required for them to react.

Combining equations 3,4 and 6, ln(1/t_i_) or lnR_p,max_ versus 1/T are expected to be linear if E_a_ is temperature independent. In the following, these were plotted, and used to obtain activation energies for the initiation step and monomer conversion respectively. These plots were also extrapolated to estimate times of inhibition and 50% monomer conversion at 4, 23, and 37 °C. These times provided estimates of when pastes might be expected to start polymerizing and solidify respectively.

#### 2.2.3 Visually observed solidification of unmixed pastes and stability of mixed paste polymerization kinetics (observed shelf life)

To visually assess paste hardening with long-term storage, double-barrel syringes containing unmixed M_10_PPG and M_10_TEG initiator and activator pastes were stored at controlled temperatures of 4, 23, and 37 °C. At 1 day, 1, 3, 6, 9 and 12 months, small portions were extruded to check for solidification. To assess stability of mixed paste polymerization kinetics at these times, for the 4 °C stored samples a portion of the composite was mixed and polymerization kinetics at 25 °C determined by FTIR-ATR (n=3).

#### 2.2.4 Polymerisation kinetics, shrinkage and heat generation of freshly prepared and mixed pastes

To compare polymerization kinetics of different mixed composites, inhibition times, maximum reaction rates and final conversions of freshly prepared and immediately mixed experimental materials were compared with those for Simplex and Cortoss. Additionally, for the experimental materials, polymerization volume shrinkage (_*φ*_) (%) and heat generation (_*ϵ*_) (kJ/cc) were calculated using the following equations.

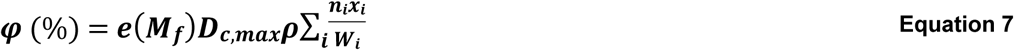

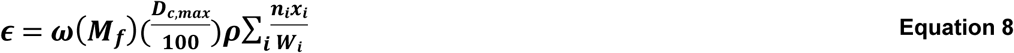

where *M_f_*, monomer mass fraction; *D_c_*, monomer conversion (%); *ρ*, composite density (g/cm^3^); _*n_i_*_; the number of C=C bonds per molecule; *W_i_* molecular weight (g/mol) of each monomer; _*x_i_*_ mass fraction of each monomer in the liquid. These assume one mole of polymerizing C=C groups gives volumetric shrinkage of 23 cm^3^/mol (_*e*_) and generates 57 kJ of heat (_***ω***_) [33].

### 2.3 Properties of set discs prepared from fresh pastes

#### 2.3.1 Polymerized disc preparation

To produce disc samples, freshly prepared and then mixed pastes were injected into metal circlips (1 mm in thickness and 10 mm in diameter). The samples were covered with an acetate sheet on top and bottom surfaces. The samples were left for 24 hr at 23 °C to allow completion of polymerization. After removal from circlips, any excess was carefully trimmed. The set samples were subsequently immersed in a tube containing 10 mL of simulated body fluid (SBF) prepared according to BS ISO 23317:2012 [34] or deionized water at 37 °C until the required test time.

#### 2.3.2 Mass and volume changes

Mass and volume changes of set composite discs after immersion in SBF at 37 °C for 0, 1, 6, 24, 48 hr and 1,2,3,4,5, 6 weeks were measured using a four-figure balance with a density kit (Mettler Toledo, Royston, UK). The percentage mass and volume change, ΔM and ΔV, were determined using the following equations [35].

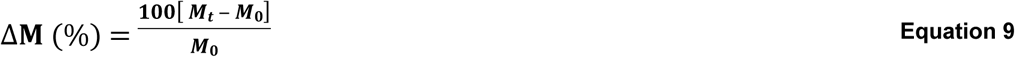

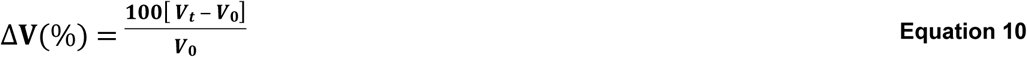

where *M*_0_ and *V*_0_ is initial mass and volume, whilst *M_t_* and *V_t_* are mass and volume at time *t* after immersion.

#### 2.3.3 Surface apatite formation

To assess ability of materials to promote surface apatite formation, disc specimens were immersed in SBF and incubated at 37 °C for 1 week (n=1). They were subsequently removed and sputtered with gold-palladium using a coating machine (Polaron E5000, East Sussex, UK) for 90 s at 20 mA. The specimens were examined under SEM (Phillip XL-30, Eindhoven, The Netherlands) operating with primary beam energy of 5 kV and a current of approximately 200 pA.

#### 2.3.4 Strontium (Sr^2+^) release

Sr^2+^ release was measured from experimental bone composites discs (n=3) immersed in 10 mL of deionized water. The specimens were incubated at 37 °C for 4 weeks. The storage solution was collected and replaced with a fresh solution at 24 hr, 1, 2, 3, and 4 weeks. The collected solution was mixed with 2 vol% nitric acid (1:1 volume ratio). Calibration standards containing Sr^2+^ of 1 ppb, 2.5 ppb, 10 ppb, 25 ppb, 100 ppb, 250 ppb, and 1 ppm were prepared using the ICP-multi element standard solution XVI (Certipur Reference Materials, Merck KGaA, Germany). The cumulative Sr^2+^ release was calculated using the following equation.

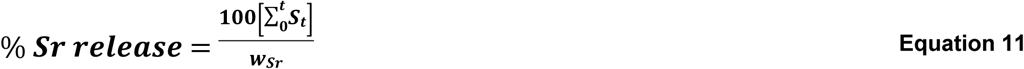

where _*wsr*_ is the initial amount of Sr^2+^ in the sample (g), *S_t_* is the amount of Sr^2+^ released into storage solution (g) collected at time *t* (hr).

#### 2.3.5 Biaxial flexural strength and fatigue life

To assess biaxial flexural strength (BFS) and fatigue performance of the materials, disc specimens were immersed in SBF and incubated at 37 °C for 4 weeks (n=25). Prior to fatigue testing, BFS of the composites was assessed using a “ball on ring” jig with a servo hydraulic testing frame (Zwick HC10, Zwick Testing Machine Ltd., Herefordshire, UK) equipped with a 1 kN load cell (n=5) [27]. The specimens’ thickness was recorded using a digital vernier calliper. The sample was placed on a ring support (8 mm in diameter). The load was applied using a 4 mm diameter spherical ball indenter at 1 mm.min^−1^ crosshead speed. The failure stress was recorded in newton (N) and the BFS (S; Pa) was calculated using the following equation [17].

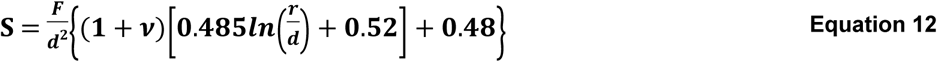

Where *F* is the load at failure (N), *d* is the specimens thickness (m), _*r*_ is the radius of circular support (m), and _*v*_ is Poison’s ratio (0.3).

For assessing fatigue performance, a sinusoidal load of 5 Hz [28] was applied to specimens using 80%, 70%, 60%, and 50% of mean BFS (n=20, 5 specimens per each level of stress). The tests were continued until fracture occurred or the requisite number of load cycles (100,000 cycles) had been applied. BFS was plotted against cycles of failure to generate classical stress-number of cycle curve (S/N curve). Failure cycle from 1^st^ to 5^th^ samples were plotted against BFS which therefore gave five S/N curves. Mean of gradient from the plots was obtained (n=5) and used to compare fatigue performance [30]. Furthermore, number of failure cycles (fatigue life) at stress level of 10 MPa was obtained by extrapolating the regression line. This value represents fatigue life of materials upon applying low stress that may be generated during normal movements such as flexion, lateral bending, or walking [36].

### 2.4 Statistical analysis

All values and errors reported throughout this study were mean (± standard deviation SD). SPSS Statistics software (version 24 for Windows, IBM, USA) was used for statistical analysis. Homogeneity of variance was assessed using Levene’s test. When variances were equal, data were analyzed using one-way analysis of variance (ANOVA) followed by post-hoc Tukey’s test for multiple comparisons. Alternatively, the Kruskal-Wallis test, followed by multiple comparison using Dunnett’s T3 tests, was used if the variances were not equal [37]. Correlation between inhibition time and final monomer conversion with storage time was tested using Pearson’s correlation. The significance value was set at *p* = 0.05. Line fitting for regression analysis was undertaken using the Linest function in Microsoft Office Excel 2016 for Windows.

Factorial analysis was used to assess the effect of MCPM level and diluent monomers on properties of composites from freshly prepared pastes. A full factorial equation for two variables each at high (10 wt% MCPM, PPGDMA) and low levels (5 wt% MCPM, TEGDMA) can be fitted using the following equation [17].

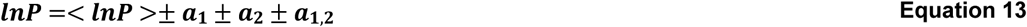

Where _*a*_1__, _*a*_2__, and _*a*_3__ were the effect of each variable on the property *P* of the composites, < *lnP* > is the average value of *lnP*. The _*a*_1,2_*, a*_1,3_ *, a*_2,3__ are interaction effects. The percentage effect of each variable, *Q*, can be calculated using the following equation.

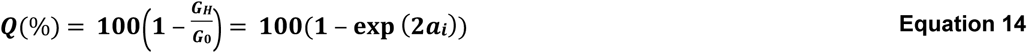

*G_H_* and *G*_0_ are the geometric average property for the samples with the variable at its high versus low value respectively. The effect of variable change was considered significant if the magnitude of _*a_i_*_ was greater than both its calculated 95% CI and interaction terms.

## 3. Results

### 3.1 FTIR studies of composite pastes

#### 3.1.1 Thermally activated polymerization of unmixed initiator paste, activation energies and predicted shelf life

An example of M_10_PPG initiator paste conversion versus time and temperature is shown in Fig 2A. Upon raising temperature from 50 to 80 °C, average t_i_ decreased from 27,000 to 350 s, whilst R_p.max_ increased from 0.010 to 0.23 %/s giving 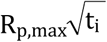 of 1.9 to 4.3 %s^-0.5^. Profiles for M_10_TEG exhibited shorter inhibition times of 1,600 to 70 s, R_p.max_ of 0.011 to 0.44 %/s and 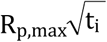 of 0.4 to 3.6 %s^-0.5^.

**Fig 2.**
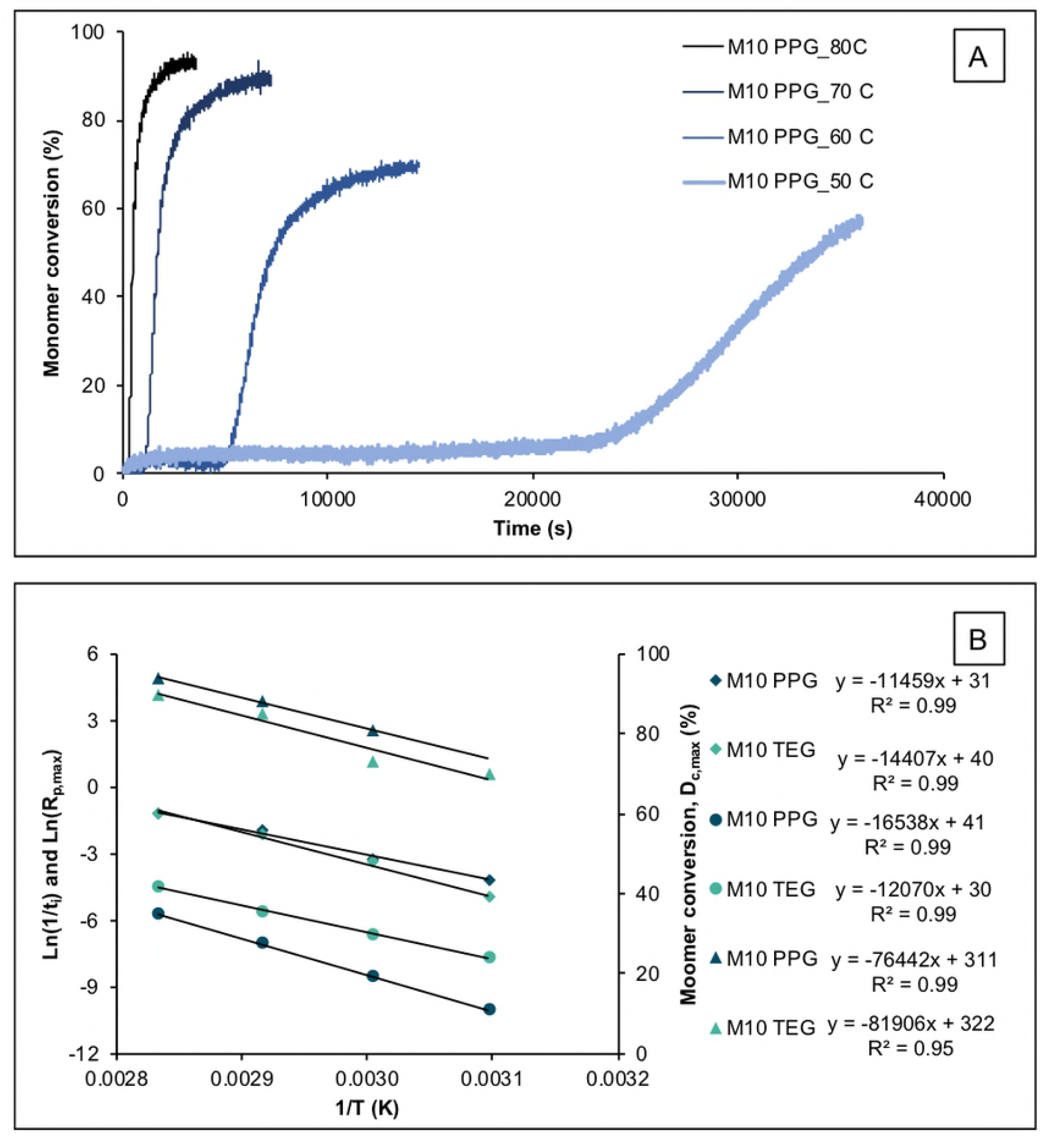
A) Example (n-1) polymerization profiles of unmixed initiator paste of M_10_PPG at different temperatures. The paste contains 3 wt% BP and 20 wt% MCPM with no activator nor TSrP. B) Average (n=1) inhibition time (t_i_; circles), R_p,max_ (diamond), and final monomer conversion (triangle) of M_10_TEG and M_10_PPG initiator pastes plotted as Ln(1/t_i_) or Ln(R_p,max_) versus inverse of temperature (Arrhenius plots). Reaction rates at a given temperature were largely similar for M_10_TEG compared with M_10_PPG although delay times and final conversions were significantly and slightly reduced respectively.

ln(1/t_i_) and ln(R_p,max_) versus inverse temperature (Fig 2B) gave linear plots (R^2^ > 0.99). M_10_PPG initiation and polymerization activation energies calculated from these were 137 and 95 kJ/mol whilst lnA terms were 41 and 31 respectively. For M_10_TEG, activation energies were 100 and 120 kJ/mol and lnA terms were 30 and 40 respectively. Extrapolation, gave M_10_PPG, t_i_ of 3 days, 1 month, and 51 months at 37, 23, and 4 °C respectively. Times for 50% conversion were comparable. Conversely, M_10_TEG t_i_ was 3 hours, 18 hours and 12 days, whilst times for 50% conversion, were 19 hours, 7 days and 7 months at 37, 23, and 4 °C respectively.

At 50% conversion, the reaction rates began to slow and tended to final conversions that declined linearly versus 1/T (Fig 2B). At 50 °C, reaction level following the 10 hours of observation was too low with M_10_PPG to enable determination of final conversion. Between 80 and 60 °C, however, final monomer conversion declined from 94 to 81% for M_10_PPG and from 90 to 73% for M_10_TEG.

#### 3.1.2 Visual solidification of unmixed pastes and stability of mixed paste polymerisation kinetics (observed shelf life)

Visual inspection indicated that, at 37 °C, both M_10_PPG and M_10_TEG initiator pastes solidified in the syringes between 1 day and 1 month. At 25 °C, M_10_PPG polymerized between 3 and 9 months whilst M_10_TEG initiator pastes solidified between 1 and 3 months. All initiator pastes stored at 4 °C, however, remained fluid even after 12 months storage.

Fig 3 shows inhibition time and monomer conversion of mixed M_10_PPG and M_10_TEG pastes that had been stored at 4 °C unmixed for up to 12 months. With M_10_PPG, inhibition time (74 s), polymerization rate (0.43 %/s), 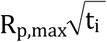 (3.7 %s^-0.5^) and final conversion (79 %) exhibited only minor change with pre-aging of pastes. With M_10_TEG, 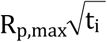 and final conversion also remained constant at 2.1 %s^-0.5^ and 70 % respectively. M_10_TEG inhibition time, however, showed a significant increase from 53 s at 24 hr to 104 s at 12 months of pre-aging (R^2^ = 0.77, *p* = 0.02).

**Fig 3.**
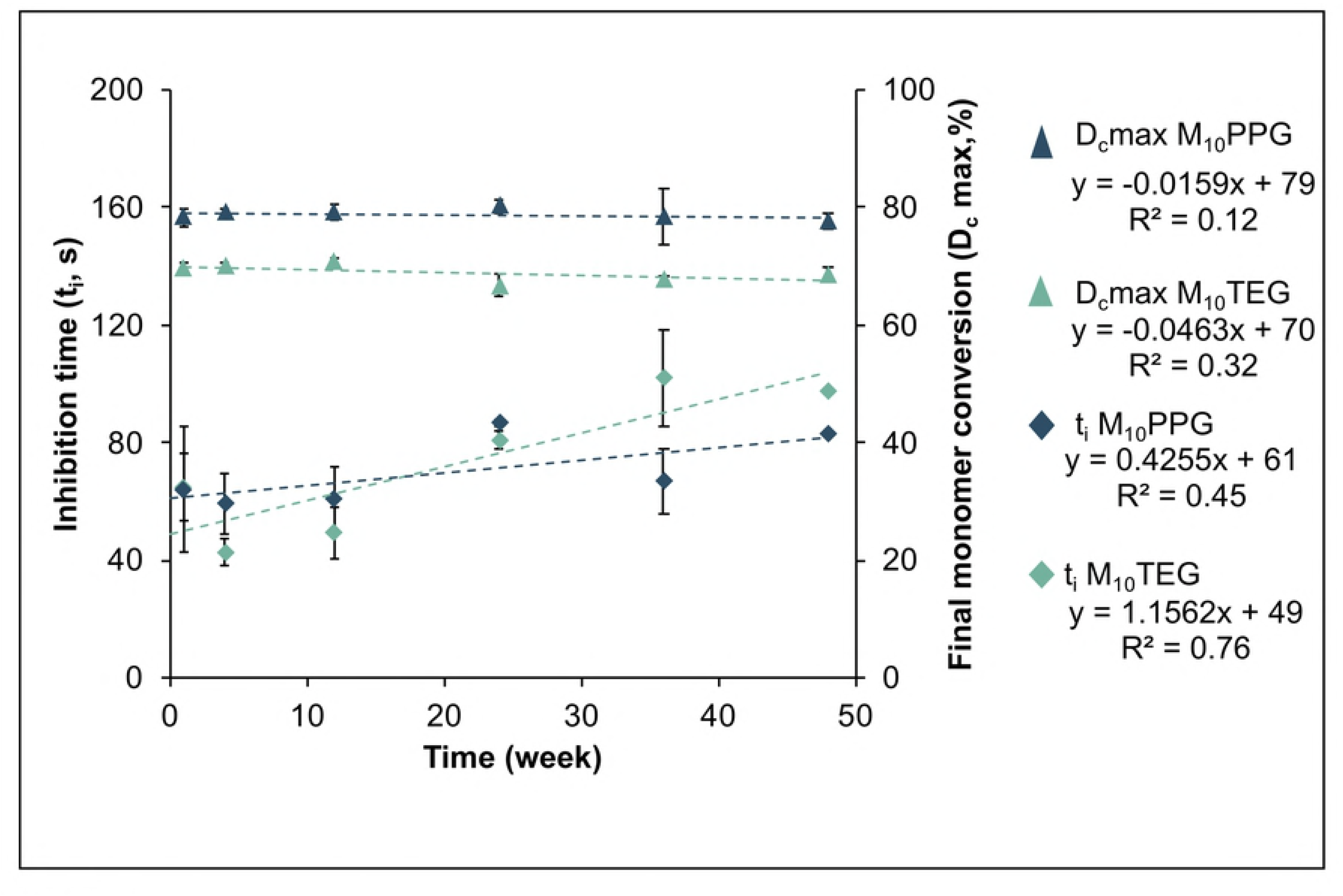
Inhibition time (diamond) and final monomer conversion (triangle) of the mixed experimental bone composites after storage unmixed at 4 °C for up to 12 months. Error bars are SD (n=3 at each time point).

#### 3.1.3 Polymerization kinetics of freshly prepared and mixed materials

Increasing MCPM level showed no significant effect on inhibition time and rate of polymerization. The shortest and longest inhibition times were observed with M_10_TEG (24 ± 4 s) and Simplex (496 ± 17 s) respectively (Fig 4-A). Inhibition times of all experimental bone composites (24 – 96 s) were shorter than that of Simplex and Cortoss (169 ± 23 s). Average inhibition time of PPGDMA based composites (85 s) was longer than that of TEGDMA based composites (24 s). Factorial analysis indicated that replacing TEGDMA by PPGDMA increased inhibition time by ~ 250 % whilst the effect of increasing MCPM level was negligible. Additionally, experimental composites exhibited comparable R_p,max_ to commercial materials (Fig 4-B). Average 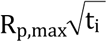 for PPGDMA, TEGDMA, Cortoss and Simplex were 4.7, 2.5, 5.4 and 10.9 %s^−0.5^ respectively.

**Fig 4.**
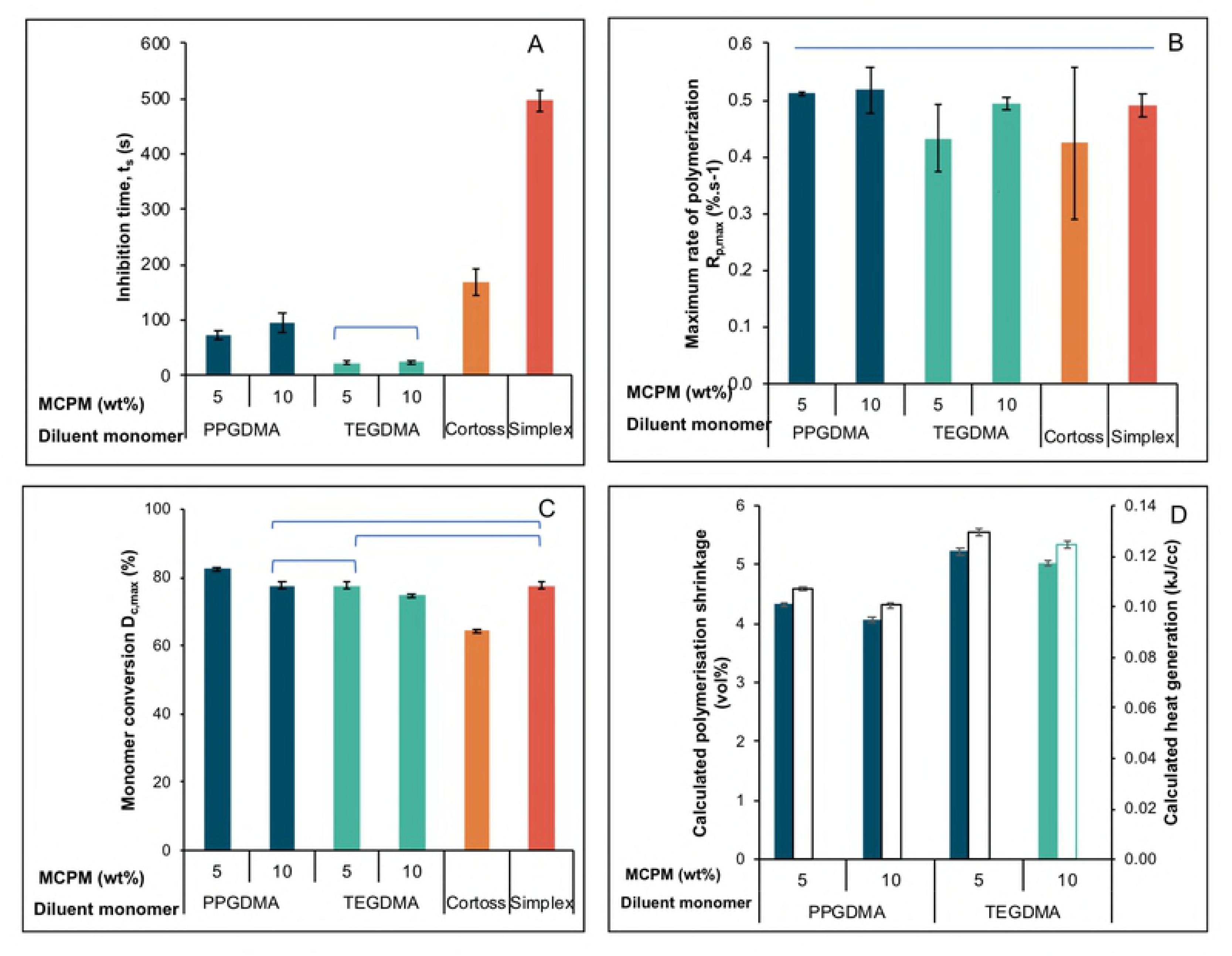
A) inhibition time, B) maximum rate of polymerization, C) final monomer conversion for experimental and commercial products, and D) calculated polymerization shrinkage and heat generation for experimental composites tested at 25 °C. Lines indicate no significant difference (*p* > 0.05). Error bars are SD (n=3).

Final monomer conversions (D_c,max_) of experimental bone composites were higher than that of Cortoss (64 ± 1 %) (Fig 4-C). M_5_PPG (82 ± 1 %) showed significantly higher final monomer conversion than Simplex (78 ± 1 %) (*p* < 0.05). Averaged final monomer conversion of PPGDMA-based bone composites (80 %) was greater than that of TEGDMA-based composites (76 %). Factorial analysis indicated that replacing TEGDMA by PPGDMA increased monomer conversion on average by 5 %. Additionally, monomer conversion of the composites was increased by 5 % upon decreasing MCPM level from 10 to 5 wt%.

Average calculated polymerization shrinkage and heat generation of TEGDMA-based experimental bone composites (5 vol% and 0.13 kJ/cc) were slightly higher than those of PPGDMA-based composites (4 vol% and 0.10 kJ/cc) (Fig 4-D). Factorial analysis indicated that the calculated shrinkage and heat generation were increased by 22% upon replacing PPGDMA by TEGDMA.

### 3.2 Mass and volume changes

Initial mass and volume change of materials increased linearly with square root of time consistent with diffusion-controlled water sorption. Simplex and Cortoss equilibrium values of mass change were 1.6 ± 0.1 wt% and 3.0 ± 0.1 wt% (Figs 5-A,B). Mass changes at later times of 3.4 ± 0.1 wt% (M_5_TEG), 4.0 ± 0.1 wt% (M_5_PPG), 4.3 ± 0.2 wt% (M_10_TEG), and 5.8 ± 0.0 wt% (M_10_PPG) were obtained for the experimental materials. Replacing TEGDMA by PPGDMA and increasing MCPM level increased mass changes at late time by 34 ± 1 % and 27 ± 3 % respectively.

**Fig 5.**
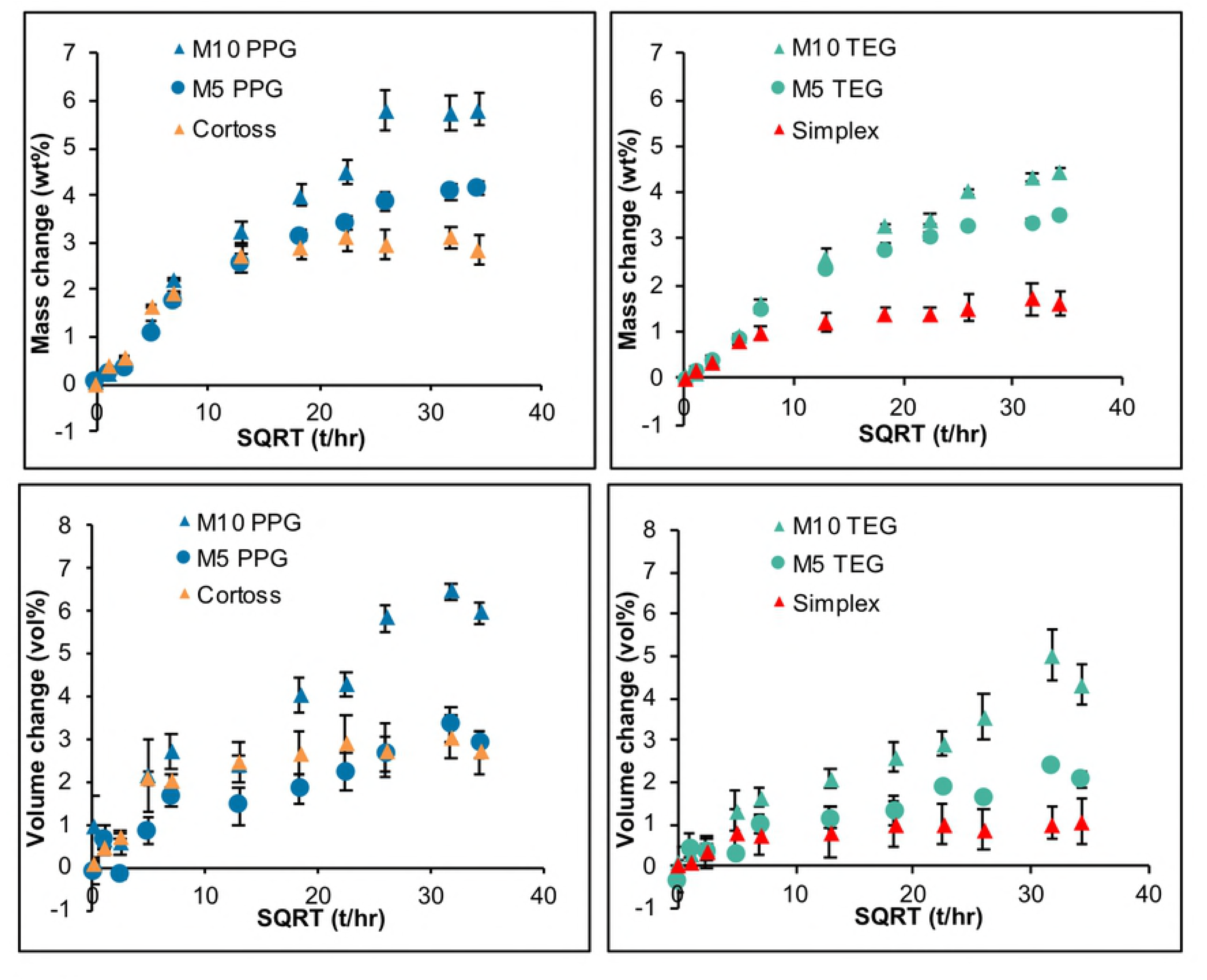
Mass and volume changes versus square root of time (hr) of all materials immersed in SBF for up to 6 weeks. Error bars are SD (n=3).

Volume change of Simplex and Cortoss reached maximum values at 1 week of 1.0 (± 0.1) and 2.8 (± 0.2) vol% (Figs 5-C,D). Later time maximum values were 1.5 (± 0.1), 3.0 (± 0.3), 4.3 (± 0.7), 6.1 (± 0.3) vol% for M_5_TEG, M_5_PPG, M_10_TEG, and M_10_PPG respectively. Factorial analysis indicated that replacing TEGDMA by PPGDMA and rising MCPM level enhanced volume change at late time by 45 ± 15 % and 109 ± 8 % respectively.

### 3.3 Surface apatite formation

At 1 week, no precipitates were observed on surfaces of M_5_TEG, Simplex, and Cortoss (Fig 6). Patchy crystals consistent with brushite were observed on some areas of M_5_PPG. Conversely, thin patchy surface apatite (~ 1 μm) layers partially covered surfaces of M_10_TEG and M_10_PPG.

**Fig 6.**
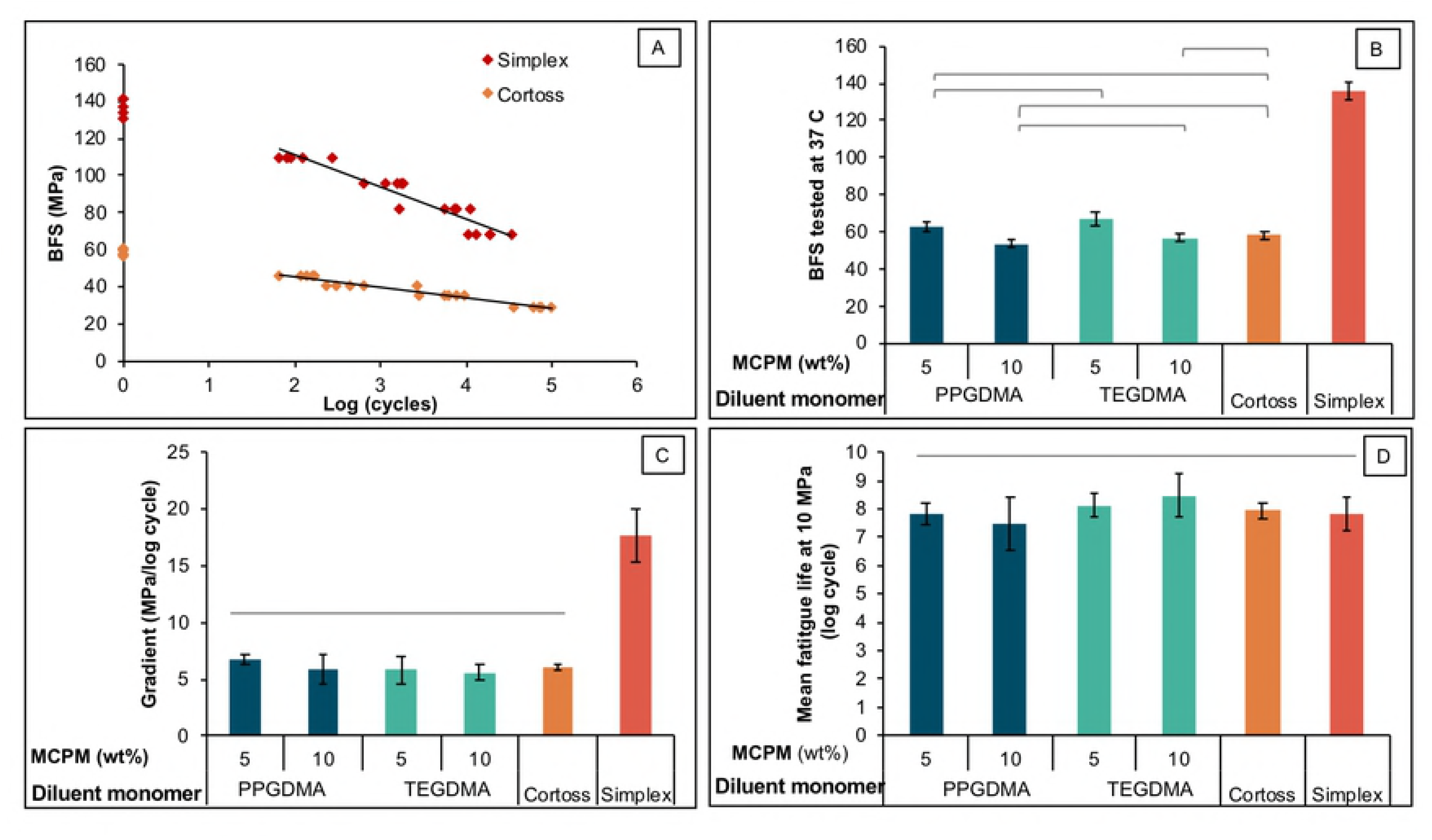
Representative SEM images for each material after immersion in SBF for up to 7 days.

### 3.4 Strontium release

The cumulative release of Sr^2+^ increased linearly with time (hr) (Fig 7). Highest and lowest rate of Sr^2+^ release was 0.0015 %.hr^−1^ and 0.0006 %.hr^−1^ observed with M_10_PPG and M_5_TEG respectively. M_10_PPG exhibited the highest accumulative Sr^2+^ release at 4 weeks (1.12 ± 0.02 %). Factorial analysis indicated that cumulative release of Sr^2+^ at 4 weeks was increased by 127 ± 14 % upon increasing MCPM level. Additionally, the release was increased by 111 ± 42 % upon replacing TEGDMA by PPGDMA.

**Fig 7.**
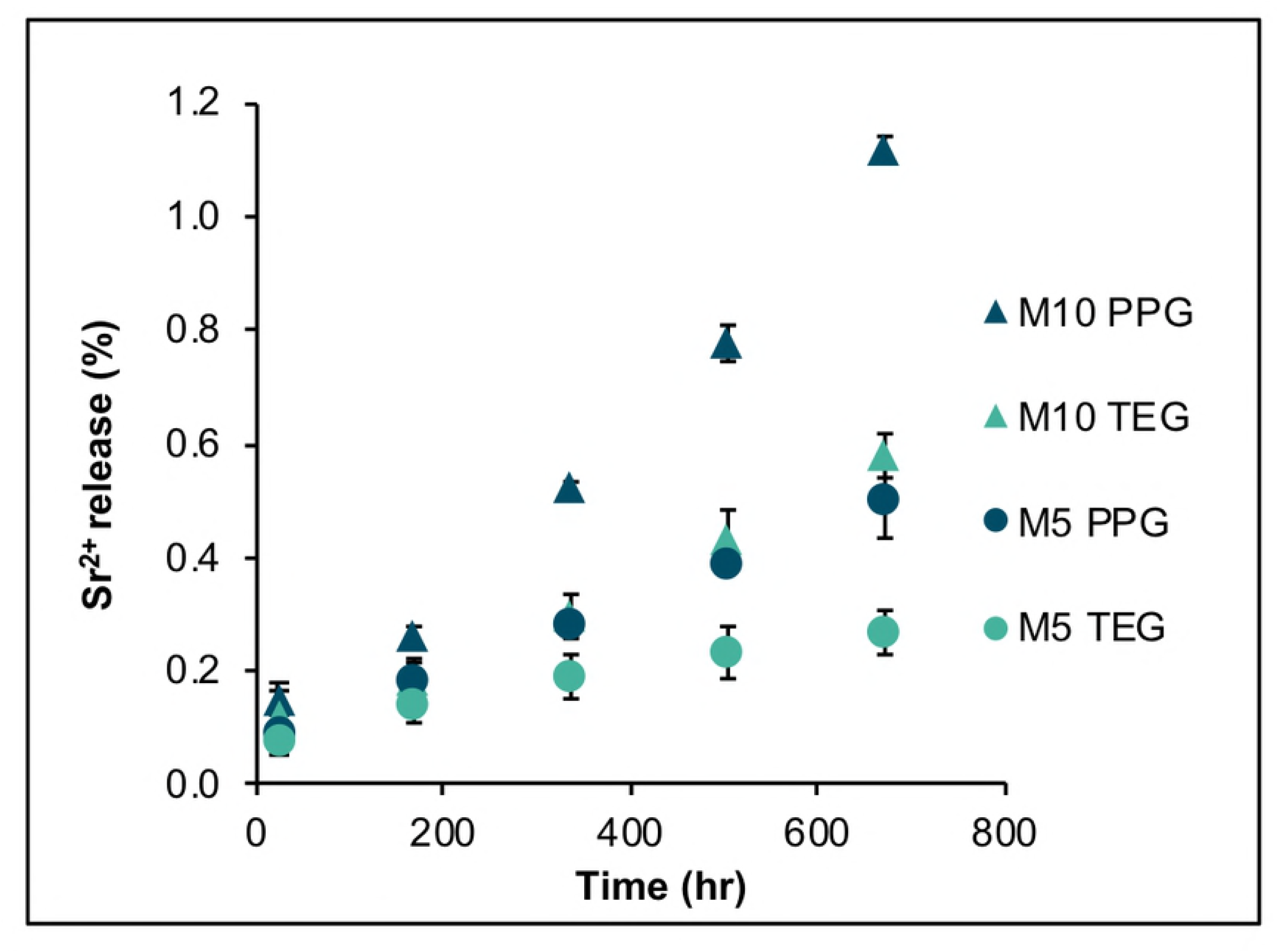
Cumulative Sr^2+^ release versus hr from bone composites immersed in deionized water up to 4 weeks. Error bars are SD (n=3).

### 3.5 Biaxial flexural strength and fatigue

The highest and lowest BFS of materials tested in SBF at controlled temperature of 37 °C was obtained from Simplex (137 ± 4 MPa) and M_10_PPG (54 ± 2 MPa) respectively (Fig 8-A). M_5_PPG had a comparable BFS (63 ± 3 MPa) to M_5_TEG (67 ± 4 MPa). The BFS of M_5_TEG was significantly higher than that of M_10_PPG (54 ± 2 MPa), M_10_TEG (57 ± 2 MPa), and Cortoss (58 ± 2 MPa). Factorial analysis showed that BFS was increased by 18 ± 5 % upon decreasing MCPM level, whilst the effect of diluent monomers was negligible.

**Fig 8.**
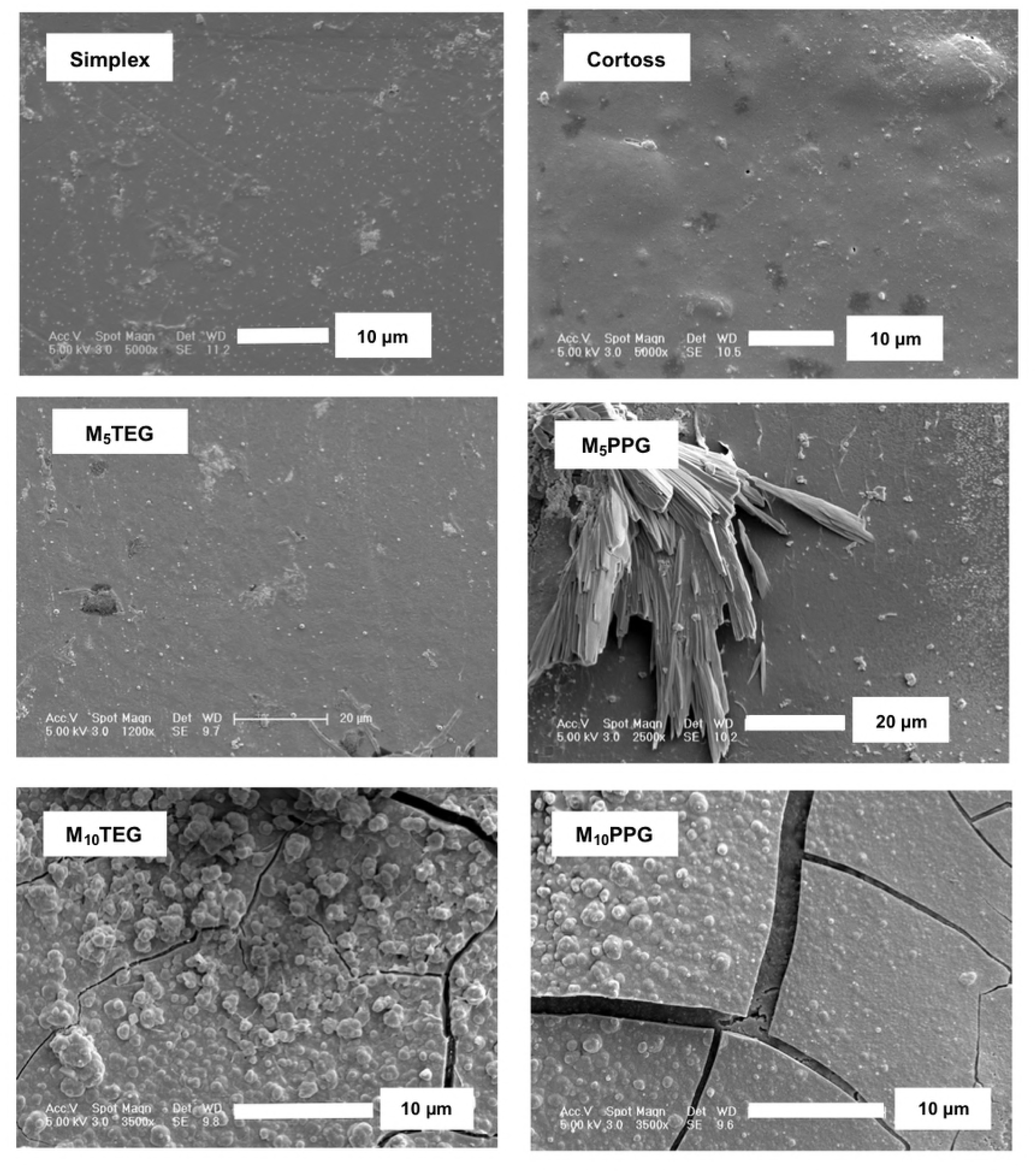
A) BFS tested in SBF at 37 °C, B) example plots of BFS versus log (cycle) (n=20), C) gradients of S/N plots in positive values for clarity purpose, and D) extrapolated fatigue life at BFS of 10 MPa. Lines indicate no significant difference (*p* > 0.05) and error bars are SD (n=5).

BFS versus logarithm of failure cycle number (S/N curve) gave straight line plots with negative gradients (Fig 8-B). The most negative gradient was observed with Simplex (−17.7 ± 2.6 MPa/log cycle) (Fig 8-C). The gradients for M_5_PPG (−6.7 ± 0.5 MPa/log cycle), M_10_PPG (−5.9 ± 1.5 MPa/log cycle), M_5_TEG (−5.9 ± 1.4 MPa/log cycle), M_10_TEG (−5.6 ± 0.8 MPa/log cycle) and Cortoss (−6.0 ± 0.3 MPa/log cycle) were comparable. Factorial analysis indicated that the effect of MCPM level and diluent monomers were negligible.

Extrapolated failure cycle values (fatigue life) at 10 MPa for the experimental composites (7.5 – 8.2 log cycle) were not significantly different from that for the commercial materials (7.8 – 7.9 log cycle) (Fig 8-D). Additionally, factorial analysis showed that MCPM level and diluent monomer had no significant effect on the fatigue life.

## 4. Discussion

This study produced bone composites that are two-paste, chemical-cured versions of previously developed Ca/Sr and PLS-containing, single paste light-cured dental composites [17]. The main change involved light activated initiator (camphorquinone) replacement with a chemical activated initiator (BP). This enables the composite to cure chemically after mixing with a separate amine activator-containing paste instead of following light activation. Additionally, however, the tertiary amine activator N,N-dimethyl-p-toluidine (DMPT) was replaced by polymerizable NTGGMA to reduce the risk of toxic amine activator leaching. Furthermore, PLR was reduced from 4:1 to 2.3:1 to enable easy mixing of the initiator and activator-containing pastes through a fine mixing tip and enhanced flow within the vertebra. The effect of MCPM level and diluent monomers on various chemical and mechanical properties were examined.

### 4.1 FTIR studies of composite pastes

The shelf life of chemically-activated bone cement is affected by storage temperature, monomer type, inhibitor, initiator and activator levels [13]. Unmixed composite paste stability is crucial to avoid premature or thermal initiated polymerization during storage or shipment. Furthermore, polymerisation kinetics following mixing must be stable and controllable to enable effective setting under clinical conditions.

In this study, a temperature-controlled FTIR-ATR system was employed to monitor polymerization kinetics. As paste at room temperature was placed on the hotter ATR plate and reaction kinetics are highly sensitive to temperature a potential error arises due to the time taken for the paste to reach the ATR temperature. This is reduced through use of thin samples. For the elevated temperature studies, this error was further minimized through ensuring inhibition times were more than 60 s. As reactions
were monitored for up to hours, this enabled a wide range of reaction temperatures and times. The reaction temperature for mixed pastes of 25 °C mimics clinical conditions before injection into the vertebra and being just slightly above room temperature minimizes temperature variability errors.

In order to understand and predict kinetics of mixed and unmixed composite paste polymerization under different conditions, reaction mechanisms and theories were employed. The mechanism for dimethacrylate reaction used in the derivation of equations 3 to 5 included initiation, inhibition, propagation, crosslinking and bimolecular termination steps [38] which may be represented by

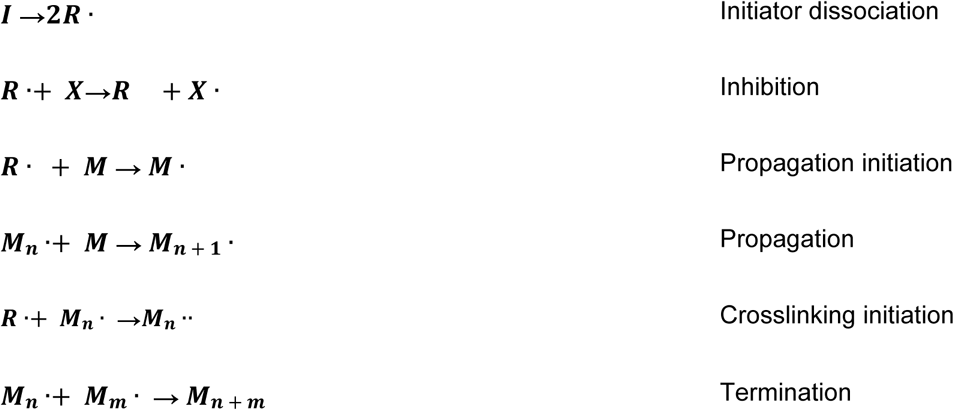

A limitation of the theory is the possibility of inhibition via routes other than by the added inhibitor. This could include free radical loss by oxygen inhibition or upon contact with surfaces such as of the filler particles or the container [39, 40]. The termination step may also occur via routes other than through bimolecular collision of two polymer free radicals [41]. Changes in relative importance of different mechanisms with reaction rate could cause errors in prediction of lower temperature stability. Lower temperature reaction rates predicted from Arrhenius plots were therefore compared with the stability and reaction kinetics of pastes that had been stored long-term.

#### 4.1.2 Paste stability and inhibition times

The observation of a delay time prior to rapid polymerization is expected from kinetic theories for both thermal and amine activated reactions [15]. For initiator and mixed pastes, this will indicate their shelf-life and time available for injection into the body (working time) respectively. According to equation 3, the inhibition time is proportional to the inhibitor concentration and inversely proportional to initiation rate. In initiator pastes, initiation rate is proportional to the benzoyl peroxide concentration. For amine activated reactions it is proportional to initiator and activator concentrations [15]. To increase initiator-paste stability and mixed paste working time, the inhibitor can therefore be increased, or the initiator and activator reduced.

Extrapolated Arrhenius plots predicted the greater low temperature stability of the PPGDMA compared with the TEGDMA initiator paste. Calculated shelf-lives, however, were ~10 times lower than those observed through long-term paste storage. This may be due to the alternative mechanisms of inhibition such as by oxygen or surface of the container when the rate of polymerization is slow. Calculated initiator paste inhibition times at 23 °C were ~10^4^ greater than those for the mixed pastes. Replacing half of the initiator by activator through paste mixing, therefore enabled rapid setting of the composite pastes.

The inhibitor in the supplied diluent TEGDMA monomer was 200 mM of MEHQ (4-methoxyphenol), whilst that of PPGDMA monomer was a mixture of 100 mM of MEHQ and 100 mM of BHT (butylated hydroxytoluene). A previous study has demonstrated that the addition of BHT enhanced the stabilisation effect of MEHQ [42] which might explain in part the observed lower inhibition times and stability of the TEGDMA initiator pastes.

The lower pre-exponential term and activation energy for the initiation step predicts faster free radical production with the TEGDMA initiator pastes at lower temperature but vice versa at high temperature. A possible explanation is that the smaller size of TEGDMA molecules reduces initial steric hindrance thereby lowering the activation energy for formation of free radicals when compared with UDMA. Conversely the larger PPGDMA molecules are of comparable size to the bulk UDMA possibly giving more comparable activation energies for free radical formation. Higher concentrations of reacting molecules but slower monomer radical formation in the PPGDMA pastes might then explain the differences in reaction kinetics.

#### 4.1.3 Polymerization rates

According to equation 4, the rate of polymerization following the inhibition period is proportional to the rate of initiation. Preferably rate of polymerization following injection of a mixed paste should be rapid to prevent leakage from the injection site or subsequent release of monomers in the body. It can potentially be raised by increasing the initiator and activator concentrations. The inhibitor may then additionally need to be raised to maintain the required working time and initiator paste stability. The rate of polymerization of the mixed pastes was comparable with that of the initiator pastes at 80 °C and ~250 times that predicted for the unmixed initiator pastes at room temperature.

From equation 5, 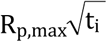 might be expected to be constant. 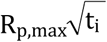 values for mixed pastes were comparable with those for the initiator pastes at the highest temperatures. A possible explanation for the decrease in 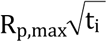 for the initiator paste with decreasing temperature, however, could be alternative free radical inhibition and termination reactions when the reaction is slow. These alternative radical removal reactions might also result in loss of the benzoyl peroxide initiator upon storage. This could then explain the increase in inhibition time of mixed pastes following long-term TEGDMA initiator paste storage.

The higher pre-exponential term for the polymerization propagation step observed with the TEGDMA initiator pastes is to be expected if higher concentration of free radicals are generated. The higher activation energy for the propagation step may be due to the TEGDMA radicals requiring more energy than UDMA or PPGDMA radicals to react with UDMA monomer.

#### 4.1.4 Maximum monomer conversions

Following 50% monomer conversion, the slowing of the dimethacrylate reaction rates can be explained by the propagation reaction that generates linear polymer chains, changing to a crosslinking process. The reaction will slow further when the conversion is sufficient to convert the material from a crosslinked rubber into a solid glassy polymer [43]. At elevated temperatures, higher conversion is required for this glass transition temperature to be reached.

Final conversions at room temperature for the PPGDMA and TEGDMA composites are comparable with values obtained using the same monomers but light activated polymerization [10]. Greater final conversion with the PPGDMA pastes could be a consequence of the longer flexible polypropylene glycol chain lowering the glass transition temperatures. Additionally, if the reaction is continuing at a fast rate when it solidifies, high concentrations of free radicals and localized heating could enable higher conversion. With the mixed fast reacting pastes, conversions at 25 °C were comparable with those achieved at 60 °C with the slower reacting initiator pastes.

#### 4.1.5 Polymerization of freshly prepared materials

Working time of PMMA bone cements that require mixing powder with liquid and transfer to a syringe for vertebroplasty should be approximately 6 to 10 min [44]. Approximately 3 minutes is required for mixing. 4 - 8 mL of PMMA cement is generally sufficient to stabilize a fractured vertebra [45]. With an injection rate of 0.15 mL / s [46], an injection time of 0.5 – 1 minutes is then required to deliver bone cement through a cannula to an affected site. This must be undertaken before the paste viscosity becomes too high for injection. This change in viscosity occurs due to swelling of the beads in the monomer phase.

For a two-paste bone composite in double-barrel syringe, the mixing takes only a few seconds. Additionally, no change in rheological properties occurs following mixing, lower volumes are required to stabilise fractures, and less heat generation compared with PMMA cement [47]. These features are a distinct advance for the composites and enable significant shortening of the required working time.

The inhibition times measured from FTIR-ATR following mixing of both the powder-liquid PMMA cement (496 s) and two-paste Cortoss bone composite (169 s) are different to final setting times cited in the literature [48] (378 s for Simplex and 345 s for Cortoss). This may be a consequence of using a different method (surface indentation), volumes of material in the test and batch number or time after production. The inhibition time of freshly prepared TEGDMA based composite was too short (23 s) indicating that more inhibitor should be included. According to equation 3, the inhibitor concentration would need to be increased 7 folds in order to bring the inhibition time up to that of Cortoss. This might additionally enhance the initiator paste shelf-life. With the PPGDMA paste, a doubling in inhibitor should give a similar inhibition time to that of Cortoss. From Equation 4, increased inhibitor should not affect the rates of polymerization. The similarities in experimental and commercial material reaction rates suggests this change would thus enable production of composites with “snap set” following sufficient working time for injection.

High final monomer conversion is required for good physical/mechanical composite properties in addition to the low risk of toxic monomer leaching [49, 50]. The final monomer conversion of Cortoss in the current study was lower than that previously obtained (80 %) using differential scanning calorimetry (DSC) [51]. DSC, however, has given higher final conversion compared to FTIR in other studies [52, 53]. Lower monomer conversion of Cortoss compared with experimental bone composites could be due to different primary base monomers. Generally, high glass transition temperature (*T*_g_) monomers give low final monomer conversion [18, 54]. Primary base monomer of Cortoss is Bis-GMA (*T_g_* = - 7.7 °C), whereas that of the experimental bone composites is UDMA (*T_g_* = - 35.3 °C) [54].

Monomer conversion of Simplex (77 %) in the current study is in good agreement with that obtained from published studies (70 %) [55, 56]. Simplex contains the monomethacrylate, methyl methacrylate (MMA), which unlike dimethacrylates, gives only linear chains and no crosslinking reaction. Consequently, complete polymerization of all methacrylate groups is required to prevent monomer leaching. Conversely, with dimethacrylate-containing composites, 50% conversion may be sufficient to bind all monomers within the resin matrix [18]. Hence 70-80% observed conversion with the experimental bone composites is expected to reduce the risk of unreacted monomer release and potentially lead to improved cytocompatibility [10].

#### 4.1.6 Polymerization heat generation and shrinkage

The lower concentration of double bonds per mole of PPGDMA contributed to the lower calculated polymerization shrinkage and heat generation of PPGDMA-based composites compared to the TEGDMA-based composites. The shrinkage of experimental bone composites in the current study was comparable to that of Cortoss (5 vol%) [57] but lower than that of PMMA bone cement (6 – 7 vol%) [58]. Additionally, as heat generation is proportional to shrinkage and a lower volume of composites is required to stabilize vertebral fractures [47], the composites should cause less thermal damage than Simplex upon placement. These properties may help to minimize gap or fibrous capsule formation and improve interfacial integrity at the bone-composite interface.

#### 4.1.7 Mass and volume changes

Mass increase due to water sorption of Simplex reached equilibrium within 1 week which is in accordance with a published study [59]. Cortoss exhibited greater mass increase compared to Simplex due probably to the lower monomer conversion, higher flexibility of polymer network, and hydrophilicity of bioactive glass contained in the composite [60].

The volume increase of Simplex in the current study (~ 2 vol %) was lower than the polymerization shrinkage reported from a published study (6 - 7 vol%) [58]. This mismatch between shrinkage and expansion may cause gap formation and induce fibrous encapsulation at the bone-cement interfaces. This poor material-bone integration may impair load transfer mechanisms leading to re-fracture or progression of cracks toward adjacent vertebra [61].

For experimental bone composites, their mass and volume changes were governed primarily by MCPM level and type of diluent monomer. Raising MCPM level enhanced water uptake leading to the increase of mass and volume as was previously observed with dental composites [18, 35]. Low crosslinking density due to the high molecular weight of PPGDMA could promote water diffusion, thereby increasing the mass and volume changes of the PPGDMA-based composites. For PPGDMA formulations, therefore, 5 to 10 wt% of MCPM was sufficient to enable hygroscopic expansion comparable with the calculated polymerization shrinkage. TEGDMA formulations, however, may requires greater than 5-10 wt% of MCPM to allow expansion to compensate polymerization shrinkage. These expansions are expected to relieve shrinkage stress and minimize gaps at composite-bone interface. This could potentially help to improve interfacial integrity and load transfer and reduce recurrent fracturing of the treated vertebra.

#### 4.1.8 Surface apatite formation

The apatite-forming ability in SBF has been adopted as a method for the determination of the bone bonding potential in biomaterials prior to any animal testing which requires large expenses and resources. It is proposed that the formation of surface apatite is associated with the ability of materials to promote *in vivo* bone bonding [62]. Other studies with MCPM-containing composites demonstrated that the level of apatite precipitation increased proportionally to time [63]. Mineral release is also expected to promote mineralization of newly formed bone [64].

When surface MCPM dissolves it disproportionates into phosphoric acid and dicalcium phosphate. Under acidic conditions, the later will precipitate as brushite. If the acid is neutralised by buffering ions in the SBF, the brushite can transform into apatite [65]. Increasing MCPM level from 5 to 10 wt% encouraged this transformation and greater
precipitation of surface apatite after immersion in SBF for 1 week. Replacing TEGDMA by PPGDMA, however, provided no obvious advantageous effect on surface apatite precipitation. In Cortoss, bioactive glass was added in an attempt to provide mineralisation and enhance bonding with bone [48]. Surface apatite was however not seen after SBF immersion for 1 week. This could be due to the slower dissolution of its calcium phosphate containing glass (combeite) [66] when compared with MCPM.

#### 4.1.9 Strontium release

Strontium release is of interest due to its potential beneficial effects for bone repair including increase of osteoblast proliferation and reduction of osteoclastic activities [22, 23, 67]. The observed linear release of Sr from experimental bone composites suggests it is not diffusion-controlled. It is possible that the level of strontium release was dependent upon its release from the surface following water enhancing polymer expansion. This would explain the increase of strontium release observed upon using PPGDMA and rising MCPM level. The results gave the effect of replacing TEGDMA by PPGDMA on Sr release as less than the effect upon doubling MCPM level. It is hypothesized that in addition to increased water sorption, MCPM may produce phosphoric acid that reacts with the tristrontium phosphate to form distrontium phosphate of higher aqueous solubility and thereby higher Sr ion release.

The release of strontium would be enhanced upon increasing surface area. A previous study showed that a bone composite provided greater interfacial stability at bone/material interface than a PMMA cement [68]. This was attributed to possibly a faster bone response, improved bone binding to mineral precipitation around the composite, and / or more effective penetration of the composite into porous bone. The greater penetration could give a large surface and encourage greater localized release of strontium to the surrounding osteoporotic vertebra. This may potentially help to increase bone mass and improve mechanical properties of the vertebra, thereby decreasing the risk of recurrent fractures. The release of strontium that increases linearly with time may also enable constant drug release which may be considered beneficial.

### 4.2 Biaxial flexural strength (BFS) and fatigue

The mean BFS values obtained from Simplex and Cortoss were consistent with those reported in a previous study [48]. Mean BFS of experimental bone composites was also comparable to that of Cortoss. Increasing hydrophilic contents and flexibility of polymer networks usually reduces strength of composites. Results from the current study showed that increasing MCPM level and replacing TEGDMA by more flexible PPGDMA had no significant effect on the strength and fatigue of the composites. This might be due to the low level of MCPM used and the enhanced monomer conversion from PPGDMA. Despite the fact that specimens were aged for 4 weeks, the mean BFS values of all experimental bone composites were greater than the 24 hr flexural strength of 50 MPa required by ISO 5833: Implants for surgery — Acrylic resin cements [69].

The highest gradient of *S*-*N* curve was observed with Simplex. This could be due to the lack of reinforcing glass fillers or glass fiber to retard crack propagation. Additionally, pores caused by the poor integration between BaSO_4_ particles and polymer matrix could also act as crack initiators [70]. It is assumed that the lower gradient of S-N curve of Cortoss and experimental bone composites compared with Simplex may result from the beneficial effects of absorbed water that could improve fatigue resistance. The water can plasticize resin matrix and increase polymer chain mobility which could enhance crack tip blunting [71]. For experimental bone composites, releasing of active ingredients may leave voids behind but the contained fibers could help to bridge the voids and slow down crack initiation [72].

In physiologic conditions, the injected bone cements are expected to penetrate through porous bone and cracks forming irregular shapes depending on the morphology of fractures [73]. Hence, the injected cement may be subjected to various stresses including torsion, flexion, and compression. A finite element analysis demonstrated that the maximum stresses generated in the injected cement after vertebroplasty may range from 5 to 15 MPa [36]. In the current study, a representative flexural stress of 10 MPa was used to extrapolate number of failure cycles thereby allowing comparison of fatigue life amongst materials. The predicted failure cycle upon applying this flexural stress of experimental composites was comparable to that commercial products (~ 10^8^ cycles). This may ensure a long-term mechanical performance of experimental bone composites.

## 5. Conclusions

Replacing diluent TEGDMA by PPGDMA provided beneficial effects such as increased inhibition time, increased final monomer conversion, and decreased calculated polymerization shrinkage and heat generation for the experimental bone composites. PPGDMA also promoted hygroscopic expansion to compensate polymerization shrinkage and enhanced strontium release. Additionally, no detrimental effect on mechanical properties of the composites was observed upon replacing TEGDMA by PPGDMA. Increasing MCPM level enhanced hygroscopic expansion, surface apatite formation, and strontium release. Increasing these reactive fillers reduced static strength of the composites but did not significantly reduce fatigue resistance of the composites.

## 6. Supporting information

S1 File. Raw data. Experimental and commercial bone composites raw data. (XLSX)

## 7. Acknowledgements

DMG has supplied monomers and fillers. SULZER supplied syringes and mixing tips. Dr. Graham Palmer, Dr. George Georgio, and Dr. Nicola Mordan provided technical support.

